# A novel histological staging of hippocampal sclerosis that is evident in grey matter loss *in vivo*

**DOI:** 10.1101/2022.12.13.520094

**Authors:** Diana Ortega-Cruz, Alicia Uceda-Heras, Juan Eugenio Iglesias, María Ascensión Zea-Sevilla, Bryan Strange, Alberto Rabano

## Abstract

**INTRODUCTION:** Hippocampal sclerosis of aging (HS) is defined by end-stage histological findings, strongly associated with limbic-predominant age-related TDP-43 encephalopathy (LATE). We aimed to characterize features of early HS to refine the understanding of its role within combined pathology.

**METHODS:** We studied 159 brain donations from the multimodal Vallecas Alzheimer’s Center Study. A staging system (0 to IV) was developed to account for HS progression and analyzed in relation to pre-mortem cognitive and MRI data.

**RESULTS:** Our HS staging system displayed a significant correlation with disease duration, cognitive performance and combined neuropathologies, especially with LATE. Two-level assessment along the hippocampal longitudinal axis revealed an anterior-posterior gradient of HS severity. *In vivo* MRI showed focally reduced hippocampal grey matter density as a function of HS staging.

**DISCUSSION:** The association of this staging system with clinical progression and structural differences supports its utility in the characterization and potential *in vivo* monitoring of HS.

## 1. Background

The human hippocampus plays a crucial role in memory encoding and retrieval, as well as attention and mood control^1,2^. As a key component of the medial temporal lobe (MTL), neuroanatomical unit is highly vulnerable to age-related pathologies, frequently associated with cognitive decline. Hippocampal sclerosis of aging (HS) is defined by severe neuronal loss and gliosis in CA1 and subiculum^1,3–7^, with CA2, CA3 and CA4 subfields apparently remaining conserved^8^. Highly advanced HS cases present a deterioration of the neuropil with intense astrocytic and microglial reaction^3–5,8,9^. Such changes result in worsening of the amnestic and cognitive symptoms of the patient^1,3–7^.

The prevalence of HS ranges from 2.8% to 30% in different autopsy series of dementia patients^4,6,10,11^. It especially affects individuals over 85 years^4,10,12^, being in fact initially described as a disease of the “oldest-old”. From a clinical perspective, amnestic symptoms associated with HS are shared with the symptomatic profile of Alzheimer’s Disease (AD). Distinguishing both profiles is challenging, such that HS is rarely diagnosed pre-mortem^13^. As a result, the definition of HS is until today limited to neuropathological observations^12^. HS can unusually occur in isolation of other neuropathologies, referred to as “pure HS”, with a frequency of 8-10% of HS cases^4,14^. Yet, HS commonly co-occurs with other pathologies, with neuronal loss in the hippocampus being disproportionate to the damage caused by these other pathologies alone.

Pathologies found to coexist with HS include those characterizing Frontotemporal Lobar Degeneration (FTLD), AD, tauopathies, Vascular Dementia and Lewy Body Disease (LBD)^3,4,6,8–10^. Yet, the association of HS is strongest with TDP-43 pathology, with 80-100% of HS cases presenting inclusions of this protein. MTL-predominant TDP-43 pathology has been extensively studied, particularly in combination with AD^15–17^. Recently, this condition was defined as limbic-predominant age-related TDP-43 encephalopathy (LATE) and tightly linked to HS^18^. Indeed, the consensus group that defined LATE suggested that both could be part of a single pathological spectrum and stratified cases into LATE HS- (without HS) and HS+ (with HS).

In parallel with its relation to LATE, other studies have suggested the existence of a precursor grade of HS (“pre-HS”) in association with TDP-43^5,10^. In this institutionalized dementia patient multimodal study, we aimed to characterize early pathological changes of HS to establish a complete staging system of the pathology. This system describes HS as a continuum, encompassing early stages not included within its definition up to now, and advanced stages as classically associated with HS. We explore the relationship between our proposed HS classification and LATE, as well as with a complete evaluation of combined neuropathology. Furthermore, we studied the association of this new staging system with cognitive progression and pre-mortem MRI structural differences, which extends its relevance towards the clinical setting.

## 2. Methods

### 2.1 Participants

This study was based on post-mortem brain donations from the Vallecas Alzheimer’s Center Study (VACS). This center is a nursing home for patients with dementia hosting a research unit (CIEN Foundation) in charge of their follow-up, including cognitive evaluation and MRI, as well as the BT-CIEN brain bank. Age at disease onset is estimated by neurologists based on medical records and interviews with caretakers. In this study, we included all brain donations (n=167) received between 2007 and 2020 from patients under follow-up during their time in the nursing home. We excluded subjects with a HS pattern consistent with epilepsy (n=1), i.e., mesial sclerosis involving predominantly CA3-4 subfields and dentate gyrus^19^. We also excluded subjects with a neuropathological diagnosis of FTLD (n=3), given its association to HS independent of aging^11^, and those whose hippocampus samples could not be evaluated due to a tumor (n=2) or extreme MTL atrophy (n=2). After applying these exclusion criteria, a total of 159 subjects were included in the study. **Table S1** provides the main demographic and clinical variables of the cohort, and **Table S2** provides the percentage of subjects presenting each component within combined dementia neuropathology. This cohort is characterized by a predominantly female population, advanced age at onset and long disease duration. Inclusion criteria of the nursing home favor particularly, though not exclusively, the admission of patients with clinical diagnosis of Alzheimer’s-type dementia, thus resulting in a high proportion of patients with high Alzheimer’s disease neuropathological change (ADNC).

### 2.2 Cognitive and functional evaluation

Routine clinical evaluation of patients in the VACS cohort includes baseline and biannual assessment of several cognitive, functional, and behavioral traits^20^. For clinicopathological analysis, we collected cognitive test results at the time of admission (baseline) and at the penultimate evaluation, to avoid the commonly observed floor effect at pre-mortem evaluation. Results from the following tests were included: semantic fluency (animal naming), severe Mini-Mental State Examination (sMMSE), *Mini Examen Cognoscitivo* (MEC) (a version of sMMSE adapted to Spanish and validated by Lobo *et al*.,^21^), Neuropsychiatric Inventory Questionnaire (NPI), Functional Assessment Staging Test (FAST), Clinical Dementia Rating including global (CDR) and partial memory (CDRm) scores, and Global Deterioration Scale (GDS).

### 2.3 Neuropathological work-up

Neuropathological procedures for brain extraction and immediate processing at BT-CIEN have been described previously^20^. Briefly, shortly after extraction, the brain is weighed and global and MTL atrophy is visually rated (0-3). The left hemisphere is fixed and cut in coronal slices for tissue block dissection. From these, 4μm sections are obtained for hematoxylin/eosin (H/E), Nissl and immunohistochemical staining (primary antibodies against amyloid-β, phospho-tau AT100, ubiquitin, total α-synuclein, and total TDP-43).

Neuropathological classification was performed according to published consensus criteria. The probability of ADNC was obtained based on Thal amyloid-β stage (Aβ), Braak tau stage (0-VI), and neuritic plaque NIA-C stage (0-3)^22^. Lewy body pathology was assessed through Braak α-synuclein staging (0-6)^23^ and Lewy Pathology Consensus criteria (LPC, 0-5)^24^. To assess cerebrovascular disease, we followed the staging defined by Deramecourt *et al*. (0-20)^25^ as well as VCING score^26^ on the vascular contribution to cognitive impairment (1-3), including separate assessment of arteriolosclerosis and cerebral amyloid angiopathy (CAA) in the occipital cortex (0-3). Occipital CAA evaluation from the VCING criteria was also applied to sections of the hippocampal body to obtain an adapted CAA measure in the hippocampus (0-3). The presence of microvascular calcifications and microinfarcts was also specifically assessed in the hippocampus. TDP-43 proteinopathy was evaluated according to the staging system (0-6) proposed by Josephs *et al*.^27^ and to LATE staging (0-3)^18^. Other pathologies assessed were argyrophilic grain disease (AGD) stage (0-3)^28^ and aging-related tau astrogliopathy (ARTAG)^29^, evaluated as present or absent.

### 2.4 Histological assessment of the hippocampus

The neuropathology of the MTL was assessed based on sections from three coronal blocks: (A) amygdala and anterior entorhinal cortex (section corresponding to levels 24-27 of the microatlas by Mai *et al*.^30^), (B) hippocampal head (levels 32-35), and (C) hippocampal body immediately posterior to the tip of the uncus (levels 39-43). Five stages of HS were defined based on qualitative evaluation of H/E histological sections from blocks (B) and (C), resulting in stages at the hippocampal head and body, respectively. Main hallmarks of these stages, especially for early HS, were specifically localized at the CA1-subiculum junction (CA1/Sub). The following operational criteria were defined for each stage:

##### Stage 0

CA1/Sub displays no disproportionate cellular changes (neither glial nor neuronal) compared to adjacent subfields, such that this junction does not stand out from CA1 nor subiculum.

##### Stage I

Evident glial hypercellularity is limited to CA1/Sub, which thus appears distinct from adjacent subfields. No disproportionate neuronal changes or cortical shrinkage are observed.

##### Stage II

Evident glial hypercellularity together with focal cortical shrinkage limited to CA1/Sub is observed. Cellular changes are comparable to Stage I, although some degree of hypertrophic astrogliosis may be present.

##### Stage III

An evident sclerotic segment is present at CA1/Sub, which, as described in the classical definition of this condition, is denoted by severe neuronal loss, astrogliosis and cortical shrinkage.

##### Stage IV

The sclerotic segment, showing severe neuronal loss, astrogliosis and cortical shrinkage, fully encompasses CA1 and at least one of its adjacent subfields, CA2 or subiculum, or most commonly both of them. Variably, this segment can further extend to CA3.

These histological hallmarks are summarized in **Figure 1** and supplementary **Table S3.** Although changes characterizing stages I, II and III may extend to neighboring regions in CA1, these did not in any case encompass the whole length of the subfield. Thus, given that CA1/Sub is key for identification of these changes, differential features at this junction are highlighted in **Figure 2**. Thus, given that CA1/Sub is key for identification of these changes, differential features at this junction are highlighted in **Figure 2**. Neuronal changes across stages are shown on Nissl staining in supplementary **Figure S1**. Subjects were classified as HS+ if any of the hippocampal head or body sections were at stage>0, and otherwise considered HS-. Similarly, among the HS+ group, if any of the regions were at stage III or IV they were classified as advanced HS, otherwise grouped as early HS (stages I and II).

**Figure 1.**
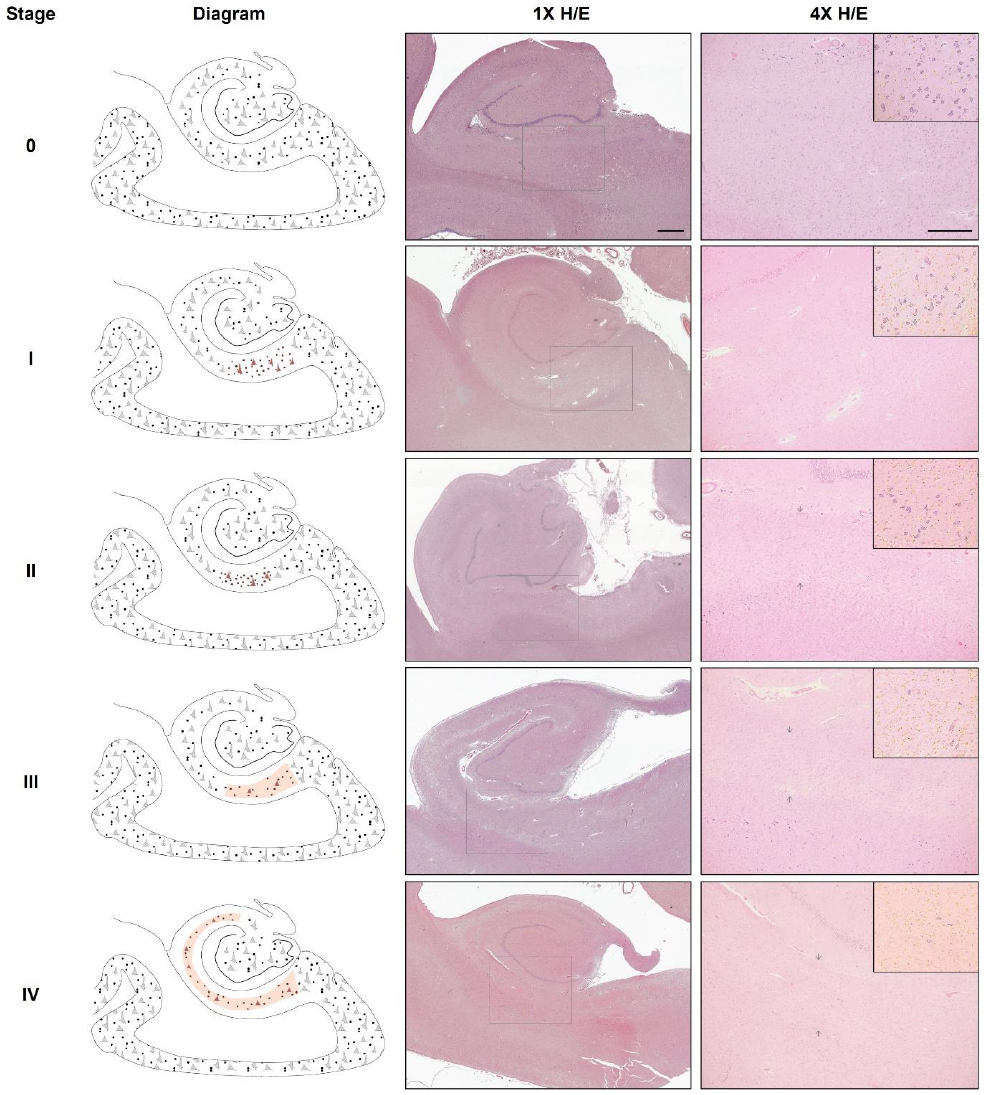
Histological features defining the HS stages within the proposed system. Left column shows schematic drawings highlighting cellular findings at each stage: stage 0 (no HS), stage I (early cellular phase), stage II (early gliotic phase), stage III (advanced short phase) and stage IV (advanced long phase). The terms “short” and “long” refer to the extent of involvement along the hippocampal transverse axis (red shaded region marks the sclerotic segment). Middle and right columns show example H/E-stained sections of hippocampal body sections at each stage, at 1x (scale bar=1mm) and 4x magnification (scale bar 4x=500μm) respectively. Square within 1x image indicates field enclosed in 4x image. Subsection in top-right corner shows outlines of neurons (dark blue) and non-neuronal cellularity (yellow) to ease their distinction in 4x image. Brightness and color saturation modified to match background color and enhance contrast. Note the shrinkage of the cortex at the CA1-subiculum junction visible in stages II, III and IV (arrows). H/E, hematoxylin/eosin.

**Figure 2.**
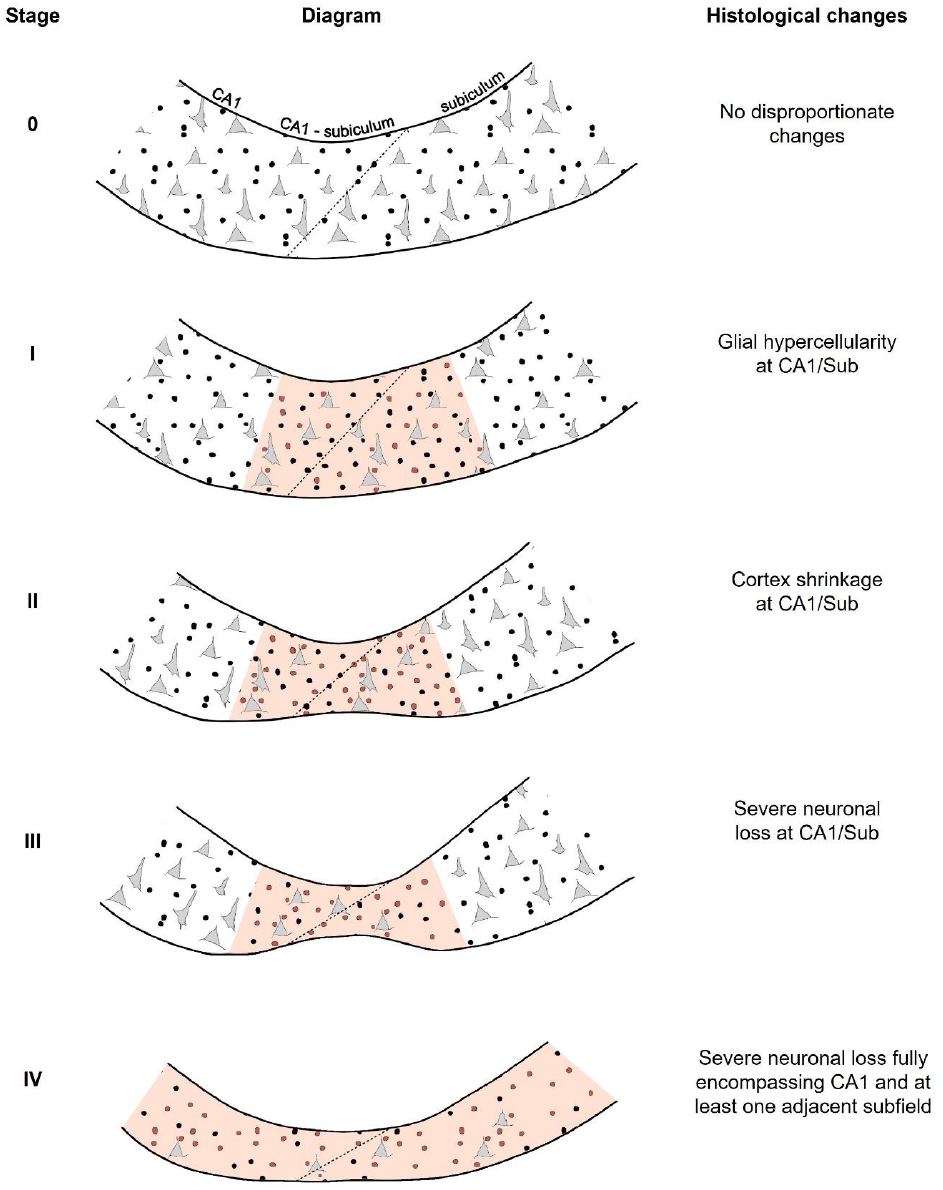
Histological hallmarks of the proposed HS staging system at the CA1-subiculum junction and adjacent regions. Schematic drawing highlighting changes in glial and neuronal cellularity as well as cortex shrinkage (present in stages II, II and IV of this system). Neurons are indicated with grey triangular shapes, and glia with black circles (with regions showing hypercellularity including red circles). The diagonal transition denoting CA1/Sub is indicated with a dotted line. The affected region that is key for distinguishing each stage is shaded in red: in stages I, II and III, the key affected region is CA1/Sub, while in stage IV sclerosis fully encompasses CA1 and at least one adjacent subfield (CA2 and CA3 are not included in this figure, shown in Fig.1). Right column shows histological changes, referring to features within a given stage that are added to those of the previous stages. At all stages, hallmarks defining each stage include those of previous stages as well. CA1/Sub, CA1-subiculum junction.

### 2.5 MRI acquisition

Among VACS subjects included in this study, 94 had at least one pre-mortem T1 MRI scan. Of these, two showed excessive motion artefacts and were discarded from analyses, leaving a final sample of 92 patients with pre-mortem T1 MRI. The scan closest to death was selected to relate to pathological evaluation, with an average antemortem interval between MRI and death of 3.02 ± 2.92 years. Additional details such as age at MRI, as well as the number of subjects included in subsequent analyses, are summarized in **Table S4**. MRI scans were acquired using a 3T MRI (Signa HDxt General Electric, Waukesha, USA) with a phased array 8 channel head coil. T1-weighted images of 1×0.469×0.469mm were obtained using a 3D sagittal sequence Fast Spoiled Gradient Recalled (FSPGR) configuration with inversion recovery.

### 2.6 Voxel-based morphometry

Voxel-based morphometry (VBM) enables the correlation of region-specific concentration of grey matter with covariates of interest^31^. This method was implemented by processing T1-weighted images using the DARTEL (Diffeomorphic Atlas Registration Tool with Exponentiated Lie Algebras) suite within SPM12 software. Briefly, images were corrected for intensity inhomogeneity, resliced to 2mm and segmented into grey matter, white matter, and CSF. Grey matter maps were spatially registered with the DARTEL technique to an in-house template, normalized to the Montreal Neurological Institute (MNI) atlas by affine transformation, modulated, and smoothed with a 6mm Gaussian kernel. Segmentations were inspected visually, and 9 subjects were discarded due to inaccurate grey matter delimitation (either at the pial or white matter boundary), resulting in 83 subjects included. Preprocessed images were compared by a voxel-wise correlation to test for differences in grey matter density (GMD) as a function of HS staging. Nuisance regressors included total intracranial volume (TIV), sex, age at scan, antemortem interval and the following neuropathological stages: Thal Aβ stage, Braak tau stage, Braak α-synuclein stage, Deramecourt vascular score, and LATE stage.

### 2.7 Hippocampal subfield segmentation

For hippocampal volume analysis, all T1-weighted scans available from 92 subjects with MRI were first processed with the FreeSurfer 7.1.1 *recon-all* stream to obtain subcortical segmentation. To increase the accuracy of subsequent steps, the subcortical segmentation was substituted by the segmentation from SynthSeg^32^ which is robust to slight movement artifacts. Automated segmentation of hippocampal subfields^33^ was then performed, and resulting labels were visually inspected to discard inaccurate segmentations. Out of 92 subjects, seven resulted in *recon-all* errors and 30 had both left and right hippocampal segmentations discarded due to incorrect hippocampus boundary delineation, resulting in 55 subjects with available volumetric MRI data.

### 2.8 Statistical analysis

RStudio 1.4.1106 was used for all statistical analyses apart from VBM. For demographic and neuropathological data, Pearson’s Chi-square tests were used to compare categorical variables between “No”, “Early” and “Advanced” HS groups. Linear models were employed to compare numeric variables if normally distributed, while the non-parametric Kruskal-Wallis test was used for non-normal numeric variables and ordinal ones. For basal and final cognitive scores, linear models were employed to assess their association with HS classification adjusting for the time interval between clinical assessment and death. Mean GMD within VBM clusters was regressed as a function of HS stages including TIV, age, sex, antemortem interval and other neuropathologies as in section 2.6, as numeric variables. The same set of independent variables was used for repeated measures ANOVAs of MRI-derived volumes.

### 2.9 Ethical approval

The BT-CIEN brain bank is an officially registered biobank by the Carlos III Research Institute (Ref: 741). BT-CIEN procedures, as well as patient follow up at the Queen Sofía Alzheimer’s Center have been approved by local health authorities (Ref: MCB/RMSFC, Expte: 12672). All brain donations were carried out under informed consent by a relative or proxy, as approved by the Ethics Committee of the Carlos III Research Institute.

## 3. Results

### 3.1 Clinical and cognitive correlation of HS staging

Our proposed staging system for HS includes early stages as part of the pathology spectrum. Therefore, subjects classified here as HS+ include those at early (I and II) and advanced stages (III and IV). To assess the clinical relevance of the proposed classification, we compared demographic and cognitive variables across groups of HS severity, dividing into subjects without HS, at early and at advanced HS stages. **Table 1** shows comparisons between these three groups as well as the correlation of HS staging to each of the included variables. Higher severity of HS was significantly more frequent in females, in spite of the strong female predominance of this cohort. In accordance with the “oldest-old” character of the pathology, HS staging and severity classification was associated with more advanced age at death, as well as longer times since diagnosis and nursing home admittance. Additionally, HS was associated with more severe atrophy, as reflected by significantly lower brain weights and higher MTL atrophy scores.

**Table 1.**
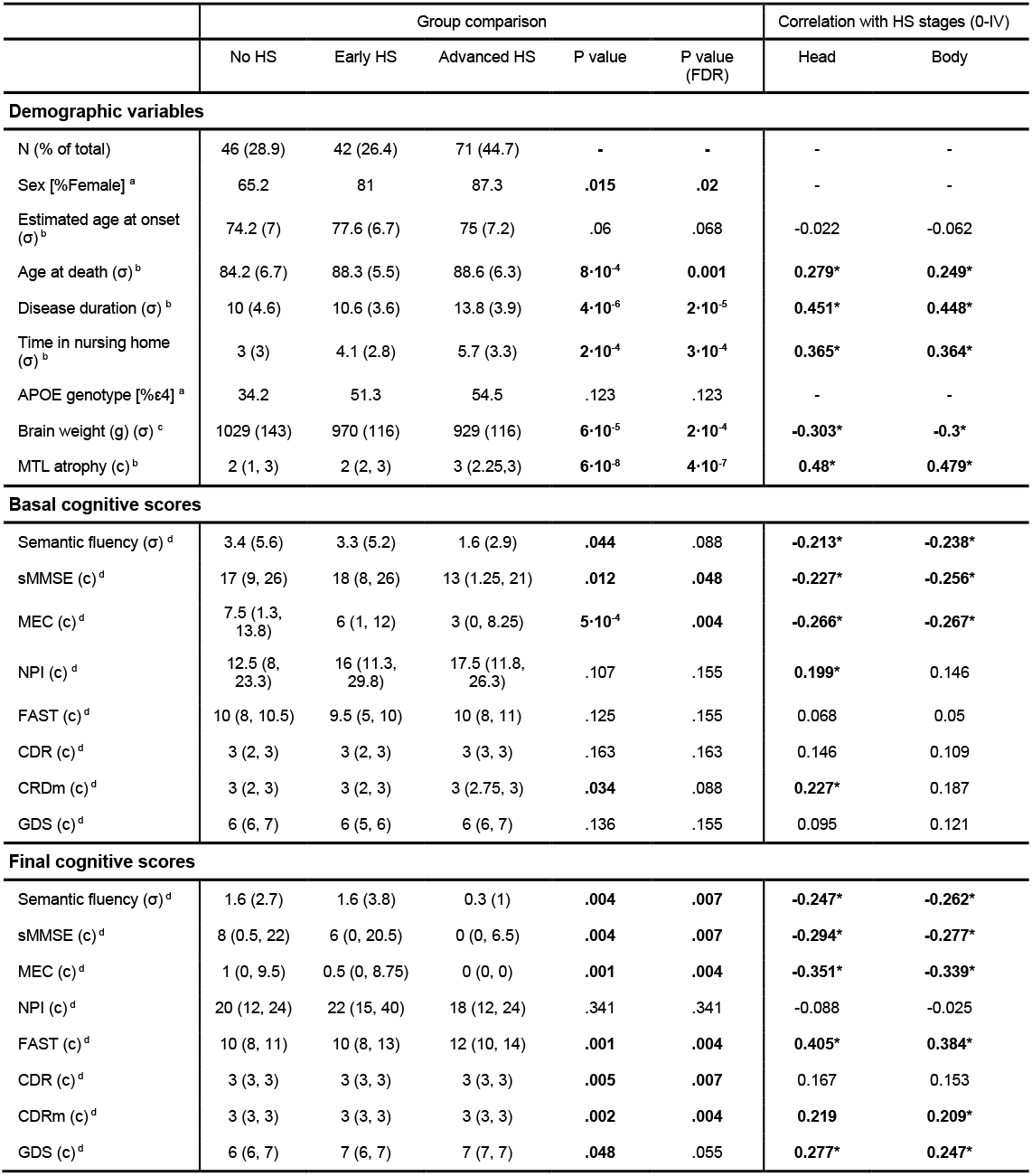
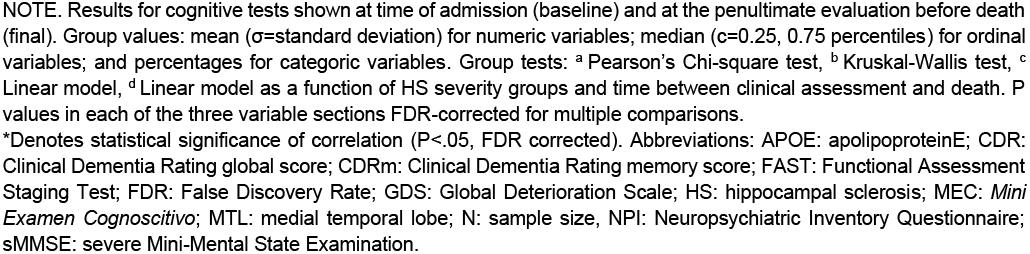
Distribution of sociodemographic, clinical, and cognitive variables across HS severity groups.

|  | Group comparison |  |  |  |  | Correlation with HS stages (0-IV) |  |
| --- | --- | --- | --- | --- | --- | --- | --- |
|  | No HS | Early HS | Advanced HS | P value | P value (FDR) | Head | Body |
| **Demographic variables** |  |  |  |  |  |  |  |
| N (% of total) | 46 (28.9) | 42 (26.4) | 71 (44.7) | - | - | - | - |
| Sex [%Female] <sup>a</sup> | 65.2 | 81 | 87.3 | **.015** | **.02** | - | - |
| Estimated age at onset (σ) <sup>b</sup> | 74.2 (7) | 77.6 (6.7) | 75 (7.2) | .06 | .068 | -0.022 | -0.062 |
| Age at death (σ) <sup>b</sup> | 84.2 (6.7) | 88.3 (5.5) | 88.6 (6.3) | **8·10<sup>-4</sup>** | **0.001** | **0.279\*** | **0.249\*** |
| Disease duration (σ) <sup>b</sup> | 10 (4.6) | 10.6 (3.6) | 13.8 (3.9) | **4·10<sup>-6</sup>** | **2·10<sup>-5</sup>** | **0.451\*** | **0.448\*** |
| Time in nursing home (σ) <sup>b</sup> | 3 (3) | 4.1 (2.8) | 5.7 (3.3) | **2·10<sup>-4</sup>** | **3·10<sup>-4</sup>** | **0.365\*** | **0.364\*** |
| APOE genotype [%ε4] <sup>a</sup> | 34.2 | 51.3 | 54.5 | .123 | .123 | - | - |
| Brain weight (g) (σ) <sup>c</sup> | 1029 (143) | 970 (116) | 929 (116) | **6·10<sup>-5</sup>** | **2·10<sup>-4</sup>** | **-0.303\*** | **-0.3\*** |
| MTL atrophy (c) <sup>b</sup> | 2 (1, 3) | 2 (2, 3) | 3 (2.25, 3) | **6·10<sup>-8</sup>** | **4·10<sup>-7</sup>** | **0.48\*** | **0.479\*** |
| **Basal cognitive scores** |  |  |  |  |  |  |  |
| Semantic fluency (σ) <sup>d</sup> | 3.4 (5.6) | 3.3 (5.2) | 1.6 (2.9) | **.044** | .088 | **-0.213\*** | **-0.238\*** |
| sMMSE (c) <sup>d</sup> | 17 (9, 26) | 18 (8, 26) | 13 (1.25, 21) | **.012** | **.048** | **-0.227\*** | **-0.256\*** |
| MEC (c) <sup>d</sup> | 7.5 (1.3, 13.8) | 6 (1, 12) | 3 (0, 8.25) | **5·10<sup>-4</sup>** | **.004** | **-0.266\*** | **-0.267\*** |
| NPI (c) <sup>d</sup> | 12.5 (8, 23.3) | 16 (11.3, 29.8) | 17.5 (11.8, 26.3) | .107 | .155 | **0.199\*** | 0.146 |
| FAST (c) <sup>d</sup> | 10 (8, 10.5) | 9.5 (5, 10) | 10 (8, 11) | .125 | .155 | 0.068 | 0.05 |
| CDR (c) <sup>d</sup> | 3 (2, 3) | 3 (2, 3) | 3 (3, 3) | .163 | .163 | 0.146 | 0.109 |
| CRDm (c) <sup>d</sup> | 3 (2, 3) | 3 (2, 3) | 3 (2.75, 3) | **.034** | .088 | **0.227\*** | 0.187 |
| GDS (c) <sup>d</sup> | 6 (6, 7) | 6 (5, 6) | 6 (6, 7) | .136 | .155 | 0.095 | 0.121 |
| **Final cognitive scores** |  |  |  |  |  |  |  |
| Semantic fluency (σ) <sup>d</sup> | 1.6 (2.7) | 1.6 (3.8) | 0.3 (1) | **.004** | **.007** | **-0.247\*** | **-0.262\*** |
| sMMSE (c) <sup>d</sup> | 8 (0.5, 22) | 6 (0, 20.5) | 0 (0, 6.5) | **.004** | **.007** | **-0.294\*** | **-0.277\*** |
| MEC (c) <sup>d</sup> | 1 (0, 9.5) | 0.5 (0, 8.75) | 0 (0, 0) | **.001** | **.004** | **-0.351\*** | **-0.339\*** |
| NPI (c) <sup>d</sup> | 20 (12, 24) | 22 (15, 40) | 18 (12, 24) | .341 | .341 | -0.088 | -0.025 |
| FAST (c) <sup>d</sup> | 10 (8, 11) | 10 (8, 13) | 12 (10, 14) | **.001** | **.004** | **0.405\*** | **0.384\*** |
| CDR (c) <sup>d</sup> | 3 (3, 3) | 3 (3, 3) | 3 (3, 3) | **.005** | **.007** | 0.167 | 0.153 |
| CDRm (c) <sup>d</sup> | 3 (3, 3) | 3 (3, 3) | 3 (3, 3) | **.002** | **.004** | **0.219** | **0.209\*** |
| GDS (c) <sup>d</sup> | 6 (6, 7) | 7 (6, 7) | 7 (7, 7) | **.048** | .055 | **0.277\*** | **0.247\*** |

NOTE. Results for cognitive tests shown at time of admission (baseline) and at the penultimate evaluation before death (final). Group values: mean ( $\sigma$ =standard deviation) for numeric variables; median (c=0.25, 0.75 percentiles) for ordinal variables; and percentages for categoric variables. Group tests: <sup>a</sup> Pearson's Chi-square test, <sup>b</sup> Kruskal-Wallis test, <sup>c</sup> Linear model, <sup>d</sup> Linear model as a function of HS severity groups and time between clinical assessment and death. P values in each of the three variable sections FDR-corrected for multiple comparisons.
\*Denotes statistical significance of correlation ( $P < .05$ , FDR corrected). Abbreviations: APOE: apolipoproteinE; CDR: Clinical Dementia Rating global score; CDRm: Clinical Dementia Rating memory score; FAST: Functional Assessment Staging Test; FDR: False Discovery Rate; GDS: Global Deterioration Scale; HS: hippocampal sclerosis; MEC: *Mini Examen Cognoscitivo*; MTL: medial temporal lobe; N: sample size, NPI: Neuropsychiatric Inventory Questionnaire; sMMSE: severe Mini-Mental State Examination.

Cognitive and functional test scores also showed significant differences between groups, particularly at the final assessment. sMMSE and MEC tests displayed significant group differences and correlation with HS staging already at the basal assessment. At the final assessment, all cognitive levels evaluated displayed significant differences across severity groups, except for neuropsychiatric NPI and GDS test scores. The specificity of these effects was confirmed by performing an additional P value correction including demographic variables, with final cognitive test differences remaining significant. Additionally, we evaluated the contribution of HS independently of LATE in linear models of cognitive loss, measured as the difference between basal and final scores, and found HS stage improves the ability of the model to explain loss in semantic fluency, sMMSE, MECT and NPI test scores (**Table S5**). Overall, our classification of HS including early and advanced stages is consistent with the expected worse cognitive performance at the final assessment close to post-mortem evaluation.

### 3.2 Anterior-posterior distribution of HS severity

HS stage was independently evaluated at two coronal levels: the hippocampal head and the anterior segment of the hippocampal body, as specified in section 2.4. For simplicity, we here refer to these two as head and body HS stages, respectively. Individual distribution of head and body stages is shown in **Figure 3A**, revealing equal stages between both regions in most cases. In subjects with different stages between regions (25.5%), HS was predominantly more advanced at the anterior portion (**Figure 3B**). The presence of more advanced stages at the hippocampal body was found in a minority of early HS cases (1.3%). Therefore, a tendency is observed towards an anterior-posterior gradient of HS severity across different stages of this system.

**Figure 3.**
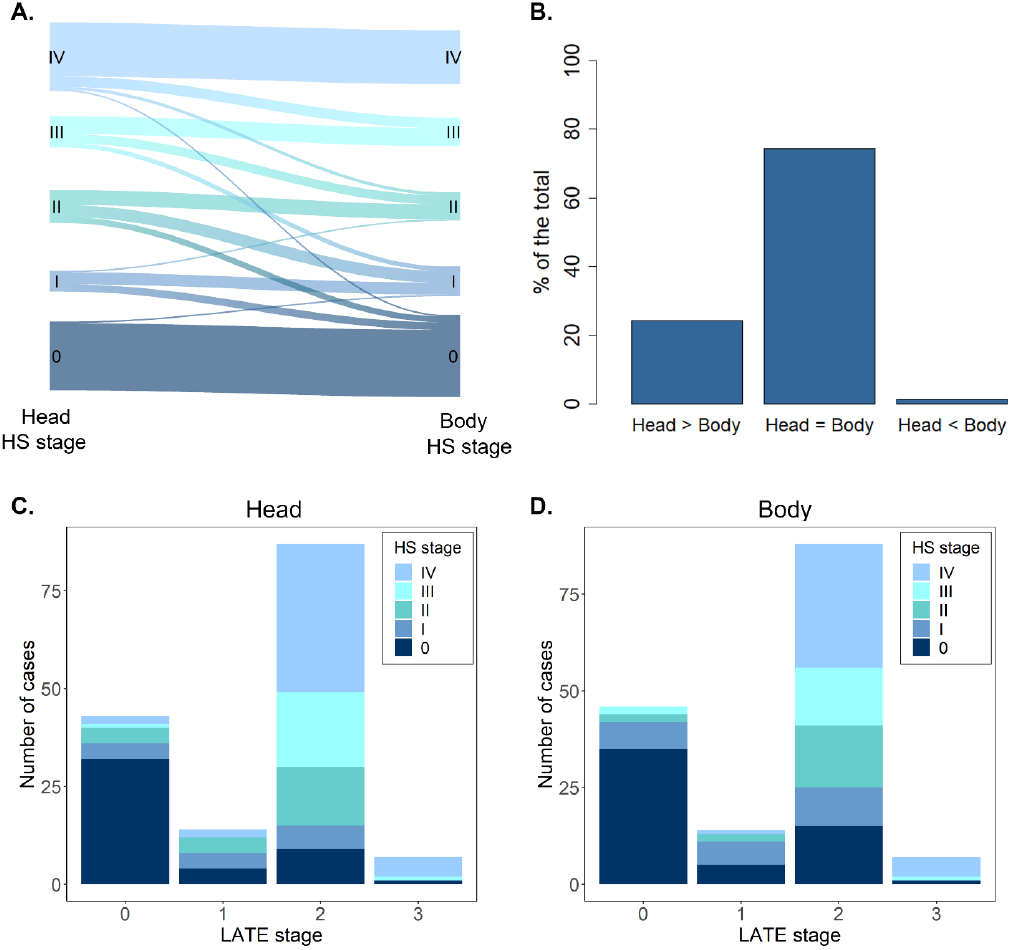
HS pathology distribution and relation to LATE. **A.** River-plot showing individual differences in HS stages between the anterior and posterior hippocampal levels assessed (head and body, respectively). **B.** Percentage of hippocampi displaying equal or different HS stages at the hippocampal head and body. **C.** Distribution of head HS stages as a function of LATE staging. **D.** Distribution of body HS stages as a function of LATE staging. HS, hippocampal sclerosis; LATE; limbic age-related TDP-43 encephalopathy.

### 3.3 Neuropathological correlation of HS staging

Since HS commonly occurs in the context of mixed pathology, a reliable staging system should present such association to other neuropathologies as well. Differences in neuropathological burden between HS severity groups as well as their correlation with HS staging are shown in **Table 2**. As expected, the proposed staging displayed the strongest correlation with TDP-43 pathology, either assessed through LATE stage or TDP-43 staging by Josephs *et al*. ^27^. Overall, 70.4% of patients presented TDP-43 inclusions at the MTL (LATE stage > 0) and 71.1% displayed either early or advanced HS changes. Such cases were not completely coincident (**Fig. 3C**), as we found HS+ subjects presenting LATE stage 0 and HS-subjects with TDP-43 inclusions (8.2% of the total in both cases). Yet, a consistent distribution between both pathologies can be identified, with subjects without TDP-43 inclusions primarily presenting HS stage 0 (20.8% of the total). Concordantly, the majority of HS+ subjects displayed LATE stage 2 (52.2% of the total), defined by the presence of TDP-43 inclusions in the hippocampus, especially those at stage IV.

**Table 2.**
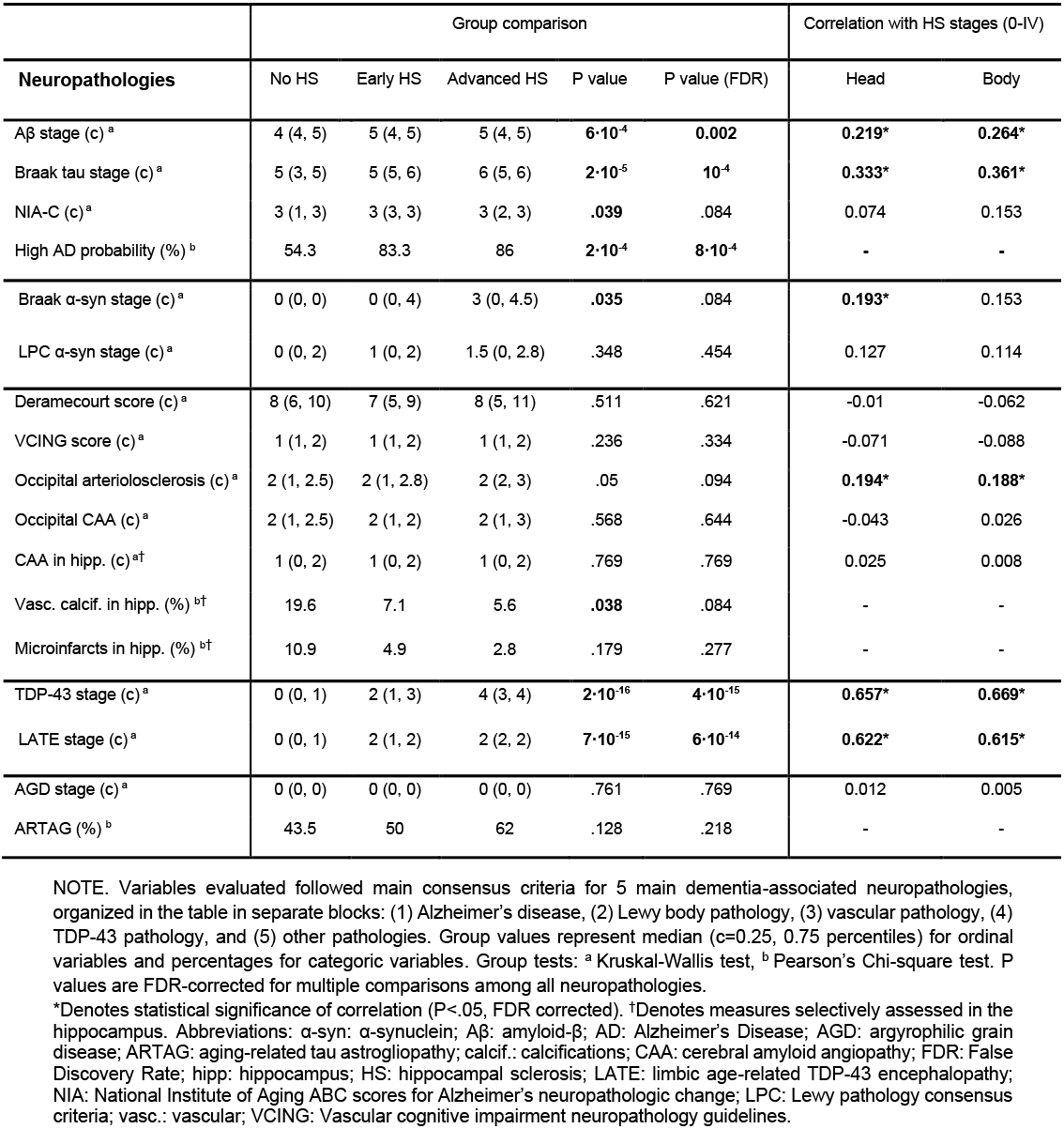
Group distribution and correlation of proposed HS staging system with other neuropathologies.

|  | Group comparison |  |  |  |  | Correlation with HS stages (0-IV) |  |
| --- | --- | --- | --- | --- | --- | --- | --- |
| **Neuropathologies** | No HS | Early HS | Advanced HS | P value | P value (FDR) | Head | Body |
| Aβ stage (c) <sup>a</sup> | 4 (4, 5) | 5 (4, 5) | 5 (4, 5) | **6·10<sup>-4</sup>** | **0.002** | **0.219\*** | **0.264\*** |
| Braak tau stage (c) <sup>a</sup> | 5 (3, 5) | 5 (5, 6) | 6 (5, 6) | **2·10<sup>-5</sup>** | **10<sup>-4</sup>** | **0.333\*** | **0.361\*** |
| NIA-C (c) <sup>a</sup> | 3 (1, 3) | 3 (3, 3) | 3 (2, 3) | **.039** | .084 | 0.074 | 0.153 |
| High AD probability (%) <sup>b</sup> | 54.3 | 83.3 | 86 | **2·10<sup>-4</sup>** | **8·10<sup>-4</sup>** | - | - |
| Braak α-syn stage (c) <sup>a</sup> | 0 (0, 0) | 0 (0, 4) | 3 (0, 4.5) | **.035** | .084 | **0.193\*** | 0.153 |
| LPC α-syn stage (c) <sup>a</sup> | 0 (0, 2) | 1 (0, 2) | 1.5 (0, 2.8) | .348 | .454 | 0.127 | 0.114 |
| Deramecourt score (c) <sup>a</sup> | 8 (6, 10) | 7 (5, 9) | 8 (5, 11) | .511 | .621 | -0.01 | -0.062 |
| VCING score (c) <sup>a</sup> | 1 (1, 2) | 1 (1, 2) | 1 (1, 2) | .236 | .334 | -0.071 | -0.088 |
| Occipital arteriolosclerosis (c) <sup>a</sup> | 2 (1, 2.5) | 2 (1, 2.8) | 2 (2, 3) | .05 | .094 | **0.194\*** | **0.188\*** |
| Occipital CAA (c) <sup>a</sup> | 2 (1, 2.5) | 2 (1, 2) | 2 (1, 3) | .568 | .644 | -0.043 | 0.026 |
| CAA in hipp. (c) <sup>a†</sup> | 1 (0, 2) | 1 (0, 2) | 1 (0, 2) | .769 | .769 | 0.025 | 0.008 |
| Vasc. calcif. in hipp. (%) <sup>b†</sup> | 19.6 | 7.1 | 5.6 | **.038** | .084 | - | - |
| Microinfarcts in hipp. (%) <sup>b†</sup> | 10.9 | 4.9 | 2.8 | .179 | .277 | - | - |
| TDP-43 stage (c) <sup>a</sup> | 0 (0, 1) | 2 (1, 3) | 4 (3, 4) | **2·10<sup>-16</sup>** | **4·10<sup>-15</sup>** | **0.657\*** | **0.669\*** |
| LATE stage (c) <sup>a</sup> | 0 (0, 1) | 2 (1, 2) | 2 (2, 2) | **7·10<sup>-15</sup>** | **6·10<sup>-14</sup>** | **0.622\*** | **0.615\*** |
| AGD stage (c) <sup>a</sup> | 0 (0, 0) | 0 (0, 0) | 0 (0, 0) | .761 | .769 | 0.012 | 0.005 |
| ARTAG (%) <sup>b</sup> | 43.5 | 50 | 62 | .128 | .218 | - | - |

NOTE. Variables evaluated followed main consensus criteria for 5 main dementia-associated neuropathologies, organized in the table in separate blocks: (1) Alzheimer's disease, (2) Lewy body pathology, (3) vascular pathology, (4) TDP-43 pathology, and (5) other pathologies. Group values represent median (c=0.25, 0.75 percentiles) for ordinal variables and percentages for categoric variables. Group tests: <sup>a</sup> Kruskal-Wallis test, <sup>b</sup> Pearson's Chi-square test. P values are FDR-corrected for multiple comparisons among all neuropathologies.
\*Denotes statistical significance of correlation (P<.05, FDR corrected). †Denotes measures selectively assessed in the hippocampus. Abbreviations: $\alpha$ -syn: α-synuclein; Aβ : amyloid-β ; AD: Alzheimer's Disease; AGD: argyrophilic grain disease; ARTAG: aging-related tau astroglipathy; calcif.: calcifications; CAA: cerebral amyloid angiopathy; FDR: False Discovery Rate; hipp: hippocampus; HS: hippocampal sclerosis; LATE: limbic age-related TDP-43 encephalopathy; NIA: National Institute of Aging ABC scores for Alzheimer's neuropathologic change; LPC: Lewy pathology consensus criteria; vasc.: vascular; VCING: Vascular cognitive impairment neuropathology guidelines.

With respect to other neuropathologies, the proposed HS classification was significantly associated with the probability of high ADNC burden, as well as with tau and Aβ stages independently. Most frequently coexisting TDP-43 and ADNC stages with HS staging are summarized in **Table S6.** Measures employed to assess global vascular burden, namely Deramecourt and VCING scores, did not show an association with HS staging (**Table 2**). Individual assessment of vascular events revealed a significant correlation of occipital arteriolosclerosis with HS stages, while that was not the case for occipital or hippocampal CAA. As for Lewy body pathology, classical Braak α-synuclein staging displayed a positive correlation with head HS stage, while differences between groups were not significant after False Discovery Rate (FDR) correction. The recently proposed LPC α-synuclein classification system displayed no correlation to HS staging. Other common pathologies found in older dementia patients such as ARTAG and AGD were not associated with HS in this series.

### 3.4 Grey matter density differences with HS staging

To further evaluate the applicability of this staging system, and its potential utility in monitoring of HS, whole-brain grey matter density from the pre-mortem MRI scan closest to death was compared among subjects with accurate grey matter segmentations available (n=83, **Table S4**). Correcting for TIV, sex, age, antemortem interval and other neuropathologies (as specified in 2.6), a focal variation of GMD relative to HS stage was found bilaterally at the hippocampus. That is, subjects at more severe HS stages at post-mortem showed reduced GMD in these regions (**Fig. 4A**). The significant cluster extended into the amygdala bilaterally and left entorhinal cortex. The average between head and body stages was taken here as a global measure of HS. **Figure 4B** shows mean GMD values within this cluster decreased linearly as a function of head and body HS stages (head: F(1,72)=83.7, P=10^−13^; body stage: F(1,72)=95.8, P=7·10^−15^). Given that significant differences were found bilaterally, we also assessed differences in mean GMD within this cluster between the left and the right hemisphere (**Figure S2**). Advanced HS stages (defined by examination of the left hemisphere) also showed reduced right GMD, with mean values being lower in the right compared to the left hippocampus across all HS stages.

**Figure 4.**
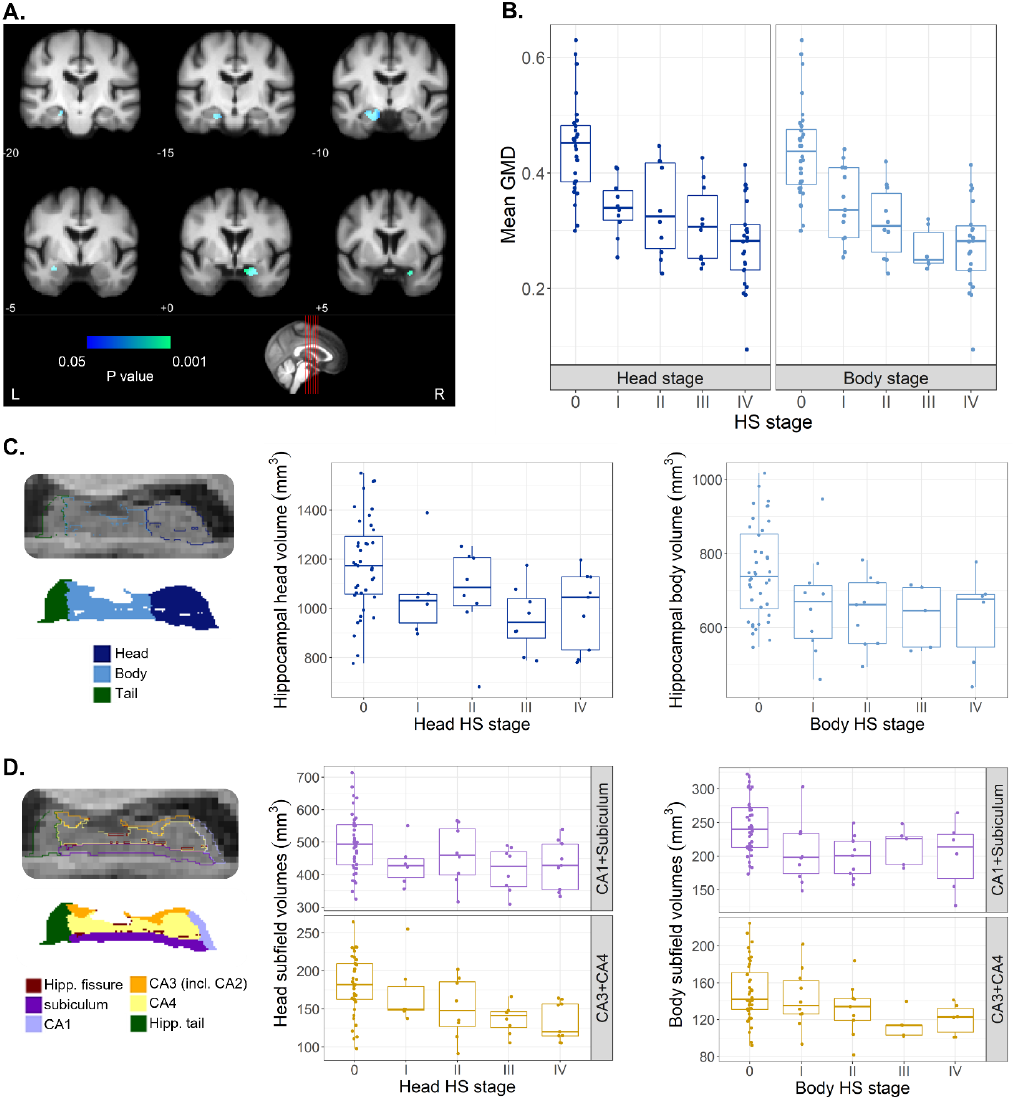
Variation of grey matter density and volume measures derived from MRI as a function HS staging. **A.** Voxel-based morphometry results of GMD as a function of the average among head and body HS stages. Effects surviving correction for multiple comparisons (Family Wise Error correction, P<.05) overlayed with an in-house template built from scans of elderly subjects, with MNI coordinates provided below each coronal section. Colormap indicates the P value of the correlation between regional-specific grey matter density and HS stage. **B.** Mean grey matter density values at the cluster of significant effects shown in A, as a function of HS stages at the head and body of the hippocampus. **C.** Volumes of hippocampal head and body as a function of respective HS stages. Head volumes displayed a significant variation as a function of HS stage. Left panel shows example segmentation labels for head, body, and tail of the hippocampus at 0.33mm resolution in sagittal view. **D.** Volume of grouped subfields (CA1+Subiculum, CA3+CA4) at hippocampal head and body, the former showing a significant variation with HS stage. Left panel shows example subfield labels resulting from segmentation at 0.33mm resolution in sagittal view. GMD, grey matter density; Hipp., hippocampal; HS, hippocampal sclerosis; incl., including; L, left; R, right.

### 3.5 Hippocampal volumetry relative to HS staging

To evaluate the relation of histologically observed subfield shrinkage with volume changes, we obtained total and subfield hippocampal volumes from automated segmentation of the T1 scan closest to death. A total of 55 subjects were included in this analysis (**Table S4**), with 40 having available data from only one hemisphere and 15 having volumes from both hemispheres. A significantly lower whole hippocampal volume was found in HS+ subjects compared to those without HS (F(1,64)=10, P=.002). To assess regional differences associated to HS, we performed a stage-dependent analysis correcting for TIV, age, sex, antemortem interval and coexisting neuropathologies. **Figure 4C** shows volumes of the head and body of the hippocampus as a function of respective HS stages. Hippocampal head volumes varied significantly as a function of HS stage (F(1,59)=8.5, P=.005) while this was not the case for the hippocampal body (F(1,59)=2.5, P=.117).

At the subfield-level, volume analysis included subiculum, CA1, CA2-3 (both segmented under the label CA3) and CA4. We grouped subiculum and CA1, as subfields classically acknowledged to be affected in HS. Similarly, we grouped CA3 and CA4 volumes. Although histological evaluation did not include CA4 due to unclear boundaries with CA3 under the optical microscope, it was included here to encompass the whole hippocampus. Estimated volumes of CA1+Subiculum and CA3+CA4 at the head and body of the hippocampus are represented in **Figure 4D**. At the hippocampal head, we found a significant relation with HS staging of CA1+Subiculum (F(1,59)=5.2, P=.026) and CA3+CA4 volumes (F(1,59)=10.7, P=.002). At the hippocampal body, volumes of both subfield groups also decreased gradually as a function of HS stage, but its contribution in this combined model was not significant.

## 4. Discussion

In this study, we have characterized early features of hippocampal sclerosis and proposed a staging system to account, for the first time, for its pathological progression. Early stages were defined by hypercellularity evidencing an increase of the glial population (stage I), followed by moderate shrinkage at the CA1-subiculum junction (stage II). Subjects at early HS stages represented one-quarter of this cohort (26.4%), an important fraction which would remain unrecognized following the classical HS definition^12^. The proposed classification of these subjects as HS+ resulted in an atypical cohort distribution, with a notably higher HS prevalence (71.1%) compared to previous studies (up to 30%) in which only advanced cases were considered^6,10,11^. In addition to these differing criteria, the advanced age and female predominance that characterize this cohort, both with reported association with HS^5,34^, contribute to explain such high prevalence. Early HS changes had been previously observed based on the presence of TDP-43 fine neurites in 42% of HS-cases in an LBD cohort^10^. While our criteria for early HS are based on histological findings rather than TDP-43 inclusions, the strong correlation of this system with LATE stages suggests the consistency of both observations.

The pathological foundations of HS are a subject of controversy^7,8,12^. Its relation to TDP-43 inclusions has been recognized through numerous studies^4,10,12,35^. In this cohort, 79.5% of HS+ cases had TDP-43 inclusions in the hippocampus, versus 19.6% for the HS-group. The parallel distribution of HS and LATE stages supports the hypothesis that these are two independent but closely interrelated pathologies. In this sense, TDP-43 could be a relevant marker for early HS, evolving into a more widespread cortical distribution in advanced HS. However, we found a minority of HS+ cases that did not present TDP-43 pathology. Future studies in larger autopsy series could address the comparison between HS+ cases with and without LATE, to discern if these present differing neuropathological signatures.

The proposed continuum for HS was also associated with tau and Aβ pathology progression. The co-occurrence of HS with tau pathology has been reported in several autopsy series^5,34,36^. Our results highlight the relevance of assessing HS in AD post-mortem studies, since it generally aggravates atrophy and cognitive decline. Additionally, α-synuclein Braak stage displayed a weak correlation with HS staging in this cohort, a relationship observed by previous authors^34,37^ but not by others^10,12,38^. Yet, these authors used differing diagnostic criteria for Lewy Body Dementia rather than Braak staging. Regarding cerebrovascular pathology, our HS classification was not associated with global vascular measures or with CAA, but it correlated with arteriolosclerosis assessment following the VCING criteria^26^. Previous studies have employed differing criteria to evaluate vascular pathology, resulting in conflicting results on its relation with HS^5,12,34,37–40^. Despite this lack of consensus, our results support the reported association of arteriolosclerosis with HS^41,42^, as well as with LATE^43,44^. A future consensual evaluation of α-synuclein and vascular pathologies throughout several cohorts would aid the clear delimitation of their relationship with HS progression.

To account for known differences in pathology susceptibility along the hippocampal anterior-posterior axis^2^, we evaluated HS at the head and body of the hippocampus. We found highly correlated but not identical HS stages between both levels. In cases with differing stages between both regions, HS was predominantly more advanced at the head level. The anterior hippocampus has been shown to present more severe TDP-43-associated atrophy^45^, as well as tau^46^ and AGD burden^47^. Our results suggest such anterior-posterior gradient is also present in HS, and thus the level at which it is evaluated should be considered. Assessment of HS at the hippocampal head will likely provide a more accurate description of its severity. Additionally, we found a transverse HS progression, with the sclerotic segment in stage IV extending to CA2 and CA3, two subfields that are rarely considered in HS evaluation^38,40^. Other HS studies have reported neuronal loss in CA2^5,9^ and CA3-4^7,48^. We highlight the relevance of hippocampal subfields beyond CA1 and subiculum in HS assessment, which would allow confirming their relation to pathology progression.

Leveraging the extraordinary setup of the VACS cohort, which includes neuroimaging follow-up, we analyzed pre-mortem structural differences. GMD in the hippocampus was found to significantly vary as a function of HS staging. Remarkably, this contribution of HS to grey matter loss was independent of other proteinopathies that severely affect the hippocampus, such as tau, TDP-43 and α-synuclein. Additionally, these results show HS-associated grey matter loss occurs bilaterally and revealed lower mean GMD values in the right compared to the left hemisphere. Several studies found bilateral HS in approximately half of the subjects, with slight predilection for the left hemisphere in unilateral cases^9,12,13,38^. While our results do not allow inferring HS stages in the right hippocampus, this cohort displayed comparable grey matter loss between hemispheres, significantly reflecting histological changes observed in the left hemisphere. MRI-derived volume analysis showed reduced hippocampal volume as a function of HS staging, particularly at the hippocampal head. This trend suggests larger sample sizes could reveal pairwise volume differences among HS stages and enable its *in vivo* classification. In agreement with histological findings, volume effects were not restricted to CA1 and subiculum, also in line with previously reported lower CA3 and CA4 volumes in HS^49^.

The strengths of this study include a relatively large cohort with an exhaustive histopathological evaluation, representing an excellent setup to evaluate the coexistence of HS with other dementia-associated neuropathologies. Furthermore, the BT-CIEN brain bank is incorporated within the VACS facilities, reducing heterogeneity in clinical data and tissue management. Besides, HS was evaluated in two regions of the hippocampus, preventing confusions from “segmental” HS^50^ and revealing anterior-posterior differences. There are, however, some limitations to this study. First, the VACS cohort is exclusive to subjects with dementia, which limits the characterization of histological features associated with the HS continuum in less severe cognitively impaired individuals. Future studies in more diverse populations that explore histopathological features of HS as well as their neuroimaging signature will position the current staging system in a more global context. Second, neuropathological examination was restricted to the left hemisphere, compromising classification of unilateral right HS cases. Third, neuroimaging analysis entailed a long mean ante-mortem interval, which could result in an underestimation of structural effects. Fourth, there was an unequal sex distribution in this cohort, hindering the evaluation of the suggested HS predominance in females^5,34^. To confirm such predominance is independent of cohort distribution and life expectancy bias, further studies with balanced sex distributions and statistical correction for age at death are needed.

In summary, this work supports the description of HS of aging as a wider pathological spectrum than initially defined, describing early stages that had been so far unrecognized. A histology-based staging system including these early stages is proposed, showing a strongly significant relation to LATE. We also report a correlation of HS staging with disease duration, cognitive status, and progression of Alzheimer’s and Lewy-type pathologies. Despite this correlation, HS staging was associated with grey matter loss independently of other neuropathologies in pre-mortem MRI. These findings highlight the relevance of HS in old age and dementia, as well as the need to find markers for its clinical diagnosis, for which MRI can be a valuable tool.

## Acknowledgements

We wish to thank the participants of the VACS cohort and their family members, as well as the CIEN Foundation staff. In particular, we thank Paloma Ruiz and Laura Saiz for their work at BT-CIEN, Paloma Mora for data retrieval, and Diego Villalon for his statistical support. This work has been supported by the Queen Sofia Foundation and the CIEN Foundation, involved in data collection for this study. DOC was supported by a fellowship from “la Caixa” Foundation (ID 100010434), with code LCF/BQ/DR20/11790034. BA and JEI were supported by a MISTI Global Seed Fund (0000000246). JEI was supported by the ERC (Starting Grant 677697), the NIH (1RF1MH123195, 1R01AG070988, 1UM1MH130981), and ARUK (ARUK-IRG2019A-003).

---

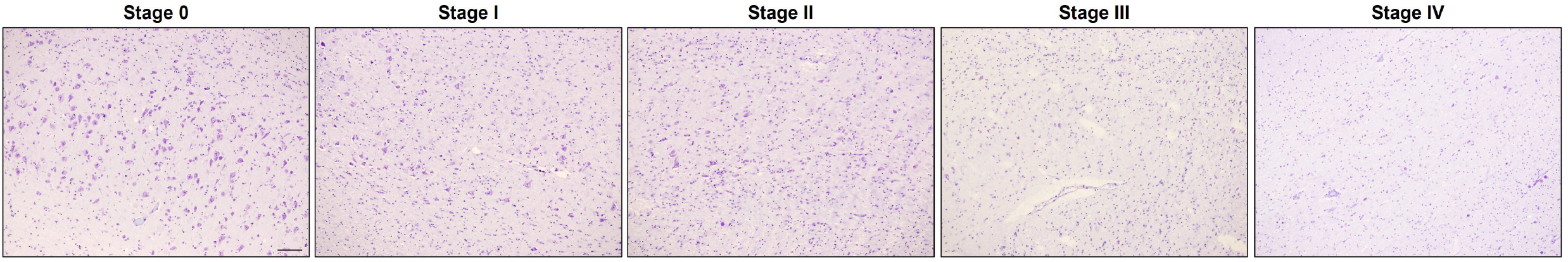

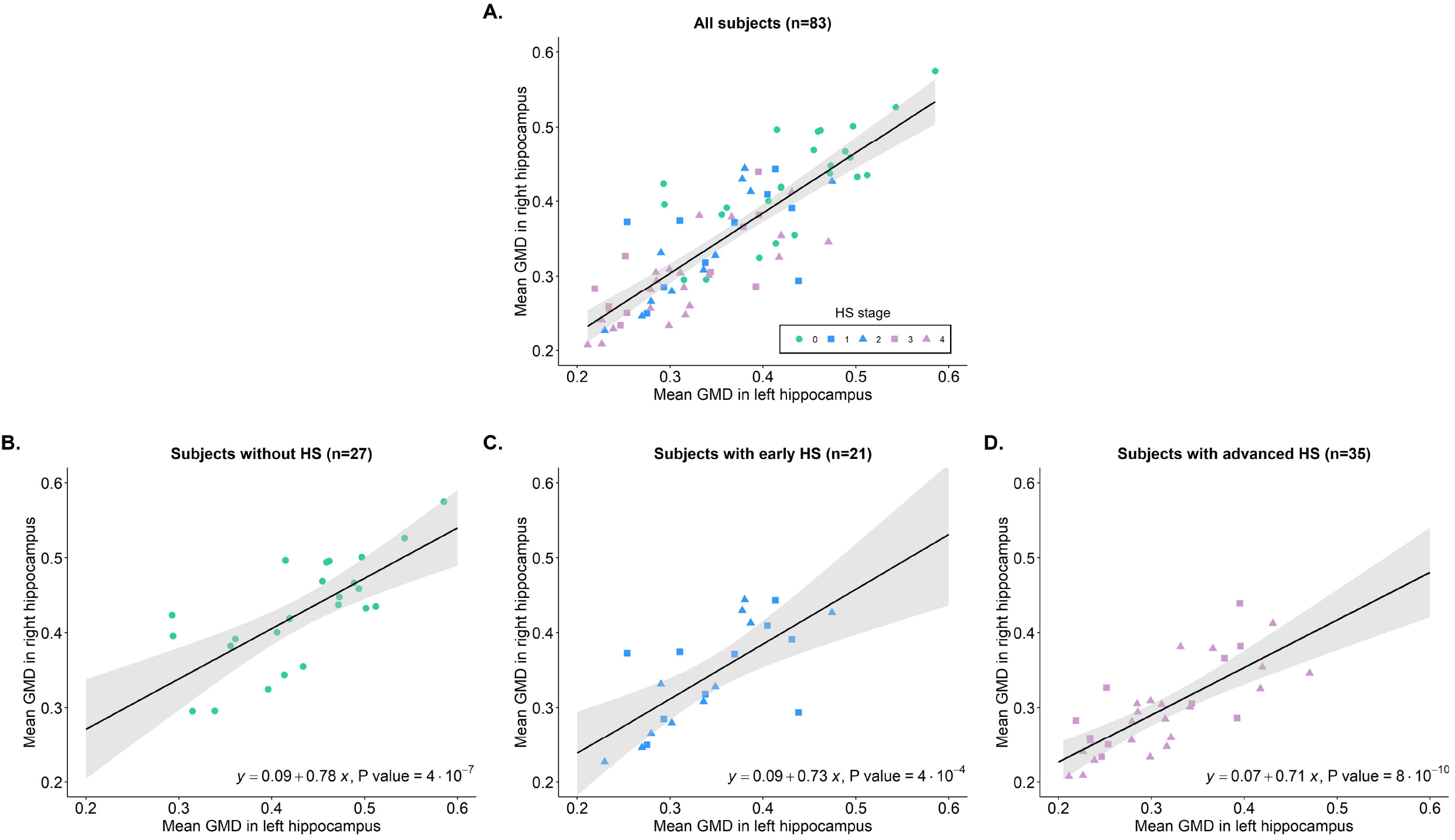

