## Supplementary Figures and Tables for "A novel histological staging of hippocampal sclerosis that is evident in grey matter loss *in vivo*"

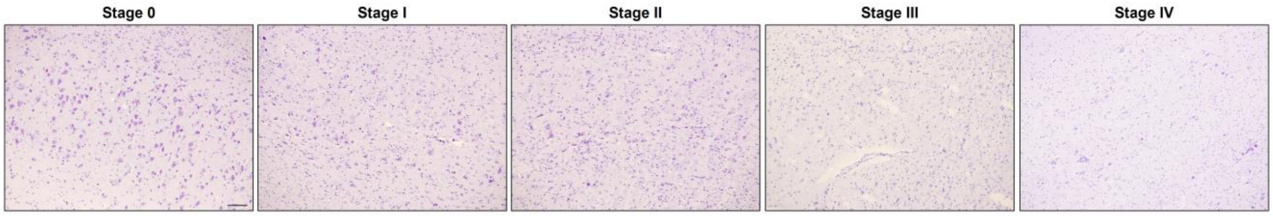

**Figure S1. Neuronal density differences between stages of the proposed system.** Nissl staining images at 10x magnification (scale bar=100μm) at the CA1-subiculum junction in example hippocampal body slides at each stage of the system. Note the severe neuronal loss detectable in advanced stages III and IV relative to stages 0, I and II.

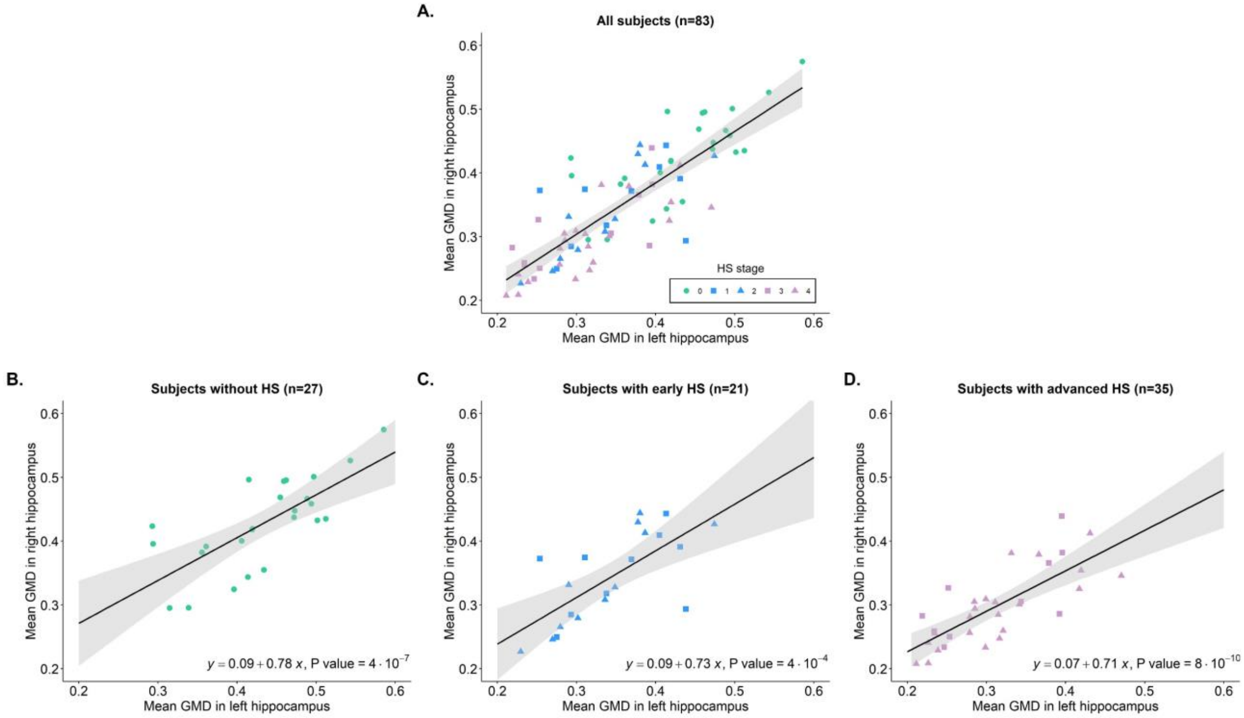

**Figure S2. GMD in the right and left hippocampus is correlated across groups of HS severity.** Mean GMD was obtained within the cluster showing significant effects in Figure 3A, combining left and right clusters to obtain regions of equal dimensions. Correlations are shown for **A.** the whole cohort, **B.** only subjects without HS, **C.** subjects at early HS stages (I and II), **D.** subjects at advanced HS stages (III and IV). For each plot, point shape and color indicates HS stage (maximum value between anterior and posterior stage). In B, C and D, equation of the fitting line and P value for the linear regression is provided. Note the slope of the fitted equation is lower than one for all groups, indicating that GMD values are lower in the right than in the left hemisphere for all stages of HS severity. Abbreviations: GMD: grey matter density. HS: hippocampal sclerosis.

| Demographics | Mean ± SD or % |
|---|---|
| Sex (%Fem) | 79.25 |
| Estimated age at onset | 75.44 ± 7.05 |
| Age at death | 87.26 ± 6.47 |
| Disease duration (years) | 11.86 ± 4.34 |
| Time in nursing home (years) | 4.46 ± 3.22 |
| Final sMMSE score | 8.25 ± 9.92 |
| APOE genotype (%ε4) | 48.25 |
| Post-mortem interval (hours) | 4.52 ± 2.14 |

**Table S1**. **Demographics and clinical variables of the studied cohort.** Abbreviations: APOE: apolipoprotein E; SD: standard deviation; sMMSE: severe Mini-Mental State Examination.

| Neuropathology burden | % |
|---|---|
| High ADNC | 76.1 |
| Intermediate ADNC | 13.8 |
| Presence of Lewy Bodies | 40.25 |
| Moderate-high vascular pathology | 36.36 |
| Presence of TDP-43 inclusions | 70.9 |
| HS (classical end-stage definition) | 44.7 |
| Early or advanced HS (proposed definition) | 71.1 |

**Table S2. Proportions of subjects of the studied cohort presenting specific neuropathologies associated with dementia.** Classification performed according to the following criteria: high and intermediate ADNC as established in reference (22) in main text; presence of Lewy Bodies determined by Braak stage>0 following reference (23); moderate-high vascular pathology following the VCING consensus criteria, reference (26); presence of TDP-43 determined by LATE stage>0 following reference (18); HS classified according to the classical end-stage definition, following reference (12), or alternatively according to the early and advanced classification proposed in this study. Abbreviations: ADNC: Alzheimer's disease neuropathological change; HS: hippocampal sclerosis; TDP-43: TAR DNA-binding protein 43.

| HS stage | | Histological change | Hippocampal subfields |
|---|---|---|---|
| None | 0 | No disproportionate changes | - |
| Early | I | Glial hypercellularity | CA1/Sub |
| | II | Cortex shrinkage | |
| Advanced | III | Severe neuronal loss | CA1/Sub |
| | IV | Severe neuronal loss | CA1, subiculum and/or CA2 |

**Table S3. Summary of histological changes detectable in specific hippocampal subfields at each stage of the proposed HS system.** "Histological changes" refers to features within a given stage that are added to those of the previous stages. Hallmarks defining each stage include those of previous stages as well (e.g., sections at stage II present cortex shrinkage together with glial hypercellularity, presented as a change in stage I). "Hippocampal subfields" refers to key regions at which these changes should be identified to classify each stage. Abbreviations: CA1/Sub: CA1-subiculum junction; HS: hippocampal sclerosis.

| **Subjects with pre-mortem MRI** | Number or Mean ± SD |
|---|---|
| Total number of subjects | 94 |
| Included in study (no excessive motion artifacts) | 92 |
| Age at MRI (included subjects) | 84.51 ± 6.44 |
| Antemortem interval (included subjects, years) | 3.02 ± 2.92 |
| **Subjects within grey matter density analysis** | |
| Number of subjects processed | 92 |
| Included in analysis (accurate GM delimitation) | 83 |
| Age at MRI | 84.28 ± 6.63 |
| Antemortem interval (years) | 3.03 ± 2.95 |
| Intracranial volume (SPM measure, $cm^3$) | 1897.51 ± 72.47 |
| **Subjects within hippocampal volumetry analysis** | |
| Number of subjects processed | 92 |
| Included in analysis (accurate subfield boundaries in at least one out of left or right hippocampus) | 55 |
| Age at MRI | 84.64 ± 6.1 |
| Antemortem interval (years) | 2.77 ± 3.11 |
| Intracranial volume (FreeSurfer measure, $cm^3$) | 1482.35 ± 152.03 |

**Table S4. Sample size and relevant parameters for all subjects with pre-mortem MRI, as well as for the subset of subjects with accurate segmentations for each analysis.** Abbreviations: GM: grey matter; MRI: Magnetic Resonance Imaging; SD: standard deviation; SPM: Statistical Parametric Mapping.

| | Models predicting difference between basal and final test scores | | |
|---|---|---|---|
| **Cognitive test** | **Model 1: LATE stage** Adjusted $R^2$ | **Model 2: LATE and HS stages** Adjusted $R^2$ | **Model 1 vs Model 2** P value |
| Semantic fluency | 0.03 | **0.087** | **.002** |
| sMMSE | 0.057 | **0.07** | **.018** |
| MEC | 0.023 | **0.067** | **.006** |
| NPI | -0.002 | **0.085** | **$2{\cdot}10^{-4}$** |
| FAST | 0.113 | 0.113 | .328 |
| CDR | **$10^{-4}$** | -0.013 | .511 |
| CDRm | 0.025 | **0.033** | .142 |
| GDS | **0.027** | 0.025 | .421 |

**Table S5. Summary of results from prediction of cognitive loss as a function of LATE and HS staging.** Cognitive loss was measured as the difference between basal and final cognitive test scores, and predicted as a function of age at death, sex, brain weight and time interval between last cognitive evaluation and death. In addition to these demographic and atrophy measures, Model 1 included LATE stage and Model 2 included both LATE and HS stages. The goodness-of-fit of each model is shown, measured by the adjusted $R^2$ parameter, with the highest value for each cognitive test shown in bold. The P value for the comparison between models 1 and 2 is shown in the right-most column of the table, with P<.05 indicating a significant difference between models (shown in bold). Abbreviations: CDR: Clinical Dementia Rating global score; CDRm: Clinical Dementia Rating memory score; FAST: Functional Assessment Staging Test; GDS: Global Deterioration Scale; HS: hippocampal sclerosis; LATE: limbic age-related TDP-43 encephalopathy; MEC: *Mini Examen Cognoscitivo*; NPI: Neuropsychiatric Inventory Questionnaire; sMMSE: severe Mini-Mental State Examination.

<table>
<tr><th colspan="2" rowspan="2">HS stages</th><th colspan="4">TDP-43 pathology stages</th><th colspan="4">ADNC stages</th></tr>
<tr><th colspan="2">LATE</th><th colspan="2">Josephs</th><th colspan="2">Braak tau</th><th colspan="2">β-amyloid</th></tr>
<tr><td>None</td><td>0</td><td>0</td><td>70%</td><td rowspan="2">0</td><td rowspan="2">30-70%</td><td>5</td><td>40%</td><td>4 – 5</td><td>35-45%</td></tr>
<tr><td rowspan="2">Early</td><td>I</td><td rowspan="2">1 – 2</td><td rowspan="2">20-55%</td><td rowspan="2">5 – 6</td><td rowspan="2">35-45%</td><td rowspan="4">5</td><td rowspan="4">70%</td></tr>
<tr><td>II</td><td>2</td><td>35%</td></tr>
<tr><td rowspan="2">Advanced</td><td>III</td><td rowspan="2">2</td><td rowspan="2">85%</td><td rowspan="2">4</td><td rowspan="2">50%</td><td rowspan="2">6</td><td rowspan="2">55%</td></tr>
<tr><td>IV</td></tr>
</table>

**Table S6. Most frequently TDP-43 and Alzheimer's disease neuropathological stages coexisting with the proposed HS staging system.** HS and TDP-43 stages display a parallel progression, while HS is mostly present in advanced stages of ADNC. Neuropathological stages for TDP-43 have been described at references (18) and (27), and AD stages in reference (22) in the main text. Most frequently coexisting TDP-43 and ADNC stages have been derived compared to the highest one among head and body HS stages. Percentages represent the proportion of cases in each HS stage (or stages, if grouped) bearing the corresponding TDP-43 or ADNC stage. Numbers are rounded (±5) to ease representation. Abbreviations: ADNC: Alzheimer's disease neuropathological change; HS: hippocampal sclerosis; LATE: limbic age-related TDP-43 encephalopathy; TDP-43: TAR DNA-binding protein 43.
